## Supplementary figures and images for "Src-dependent tyrosine-phosphorylation of NM2A has a protective role against bacterial pore-forming toxins"

### Fig S1 Revised.tif

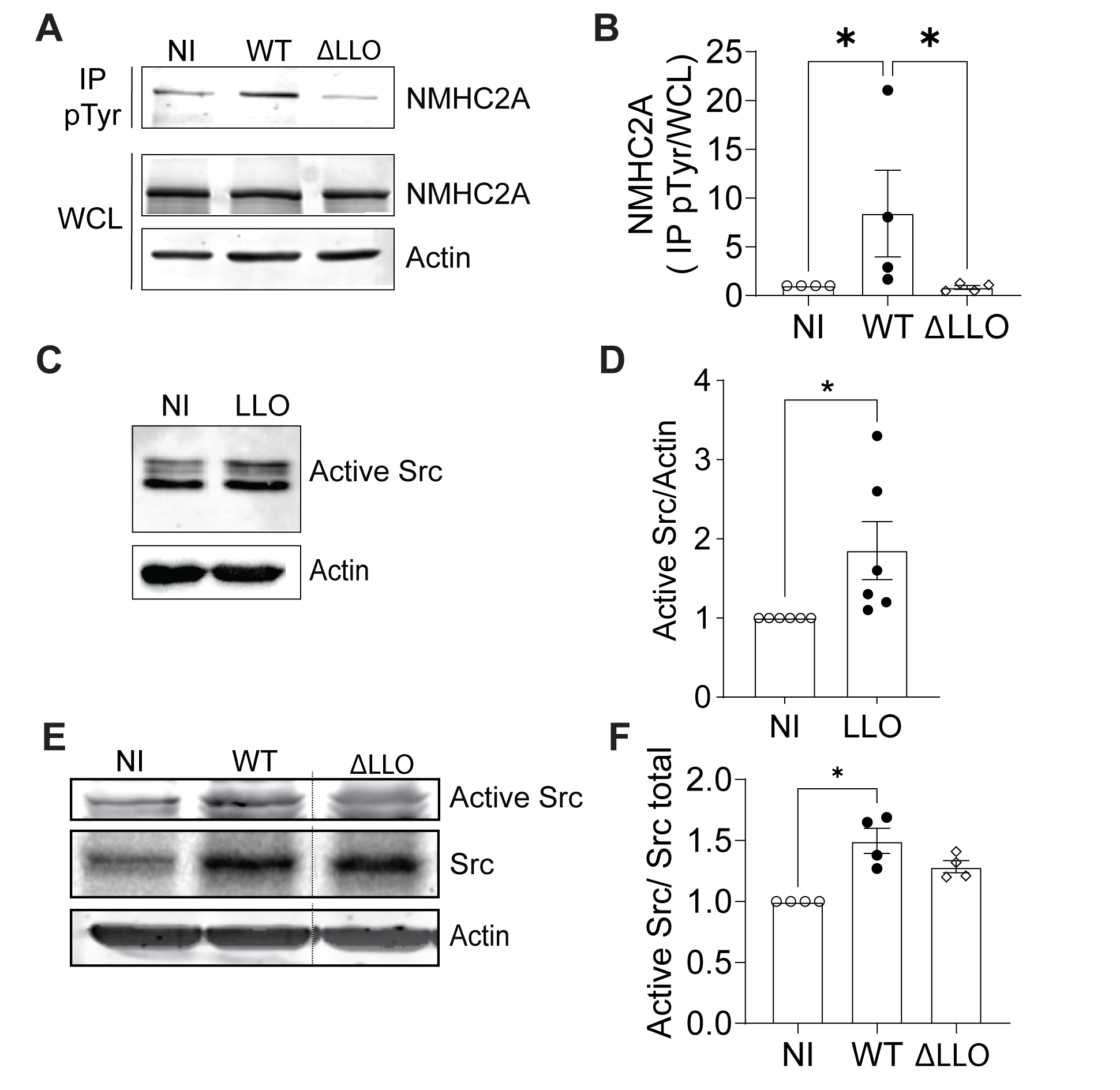

### Fig S2 Revised.tif

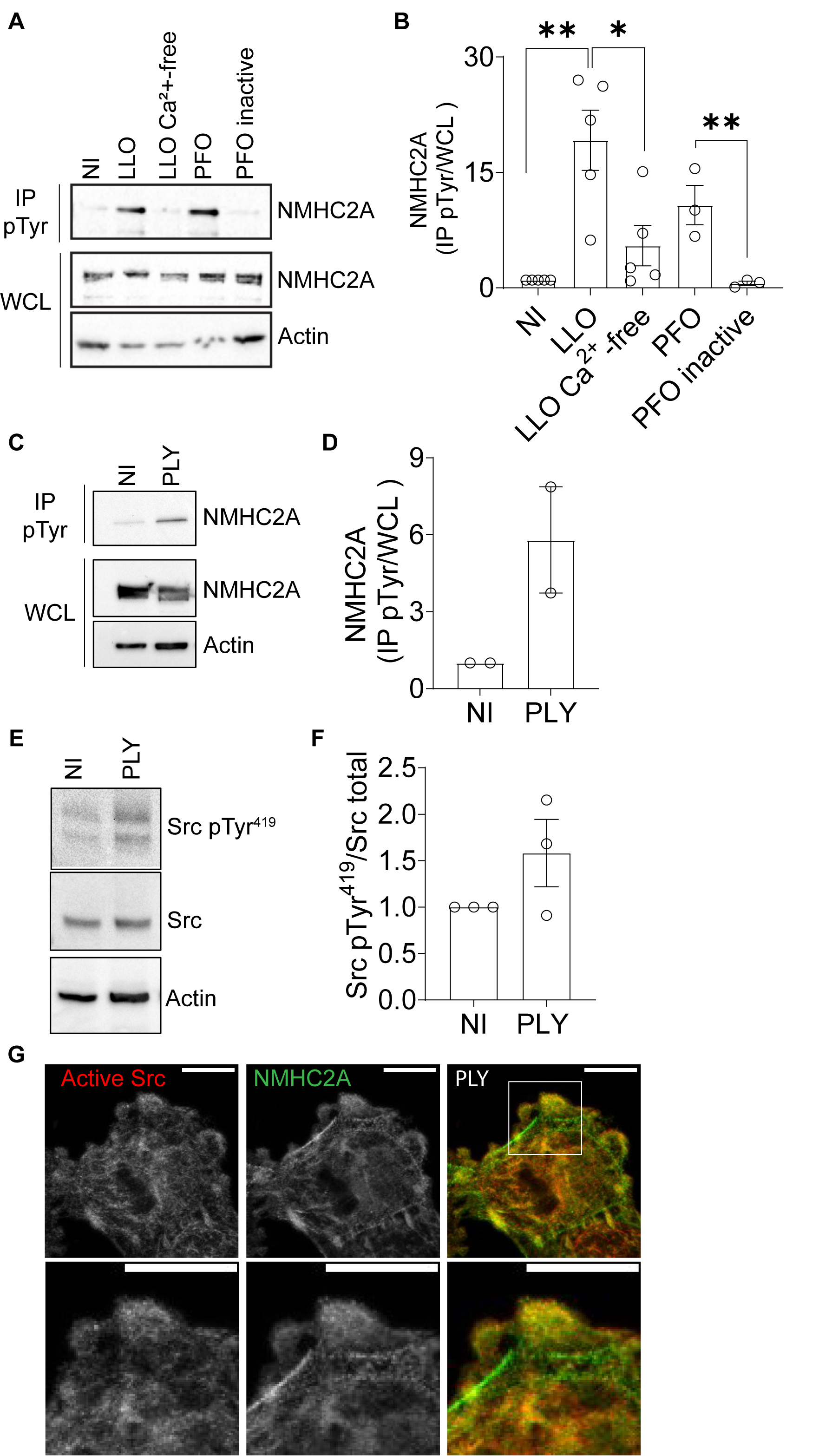

### Fig S3.tif

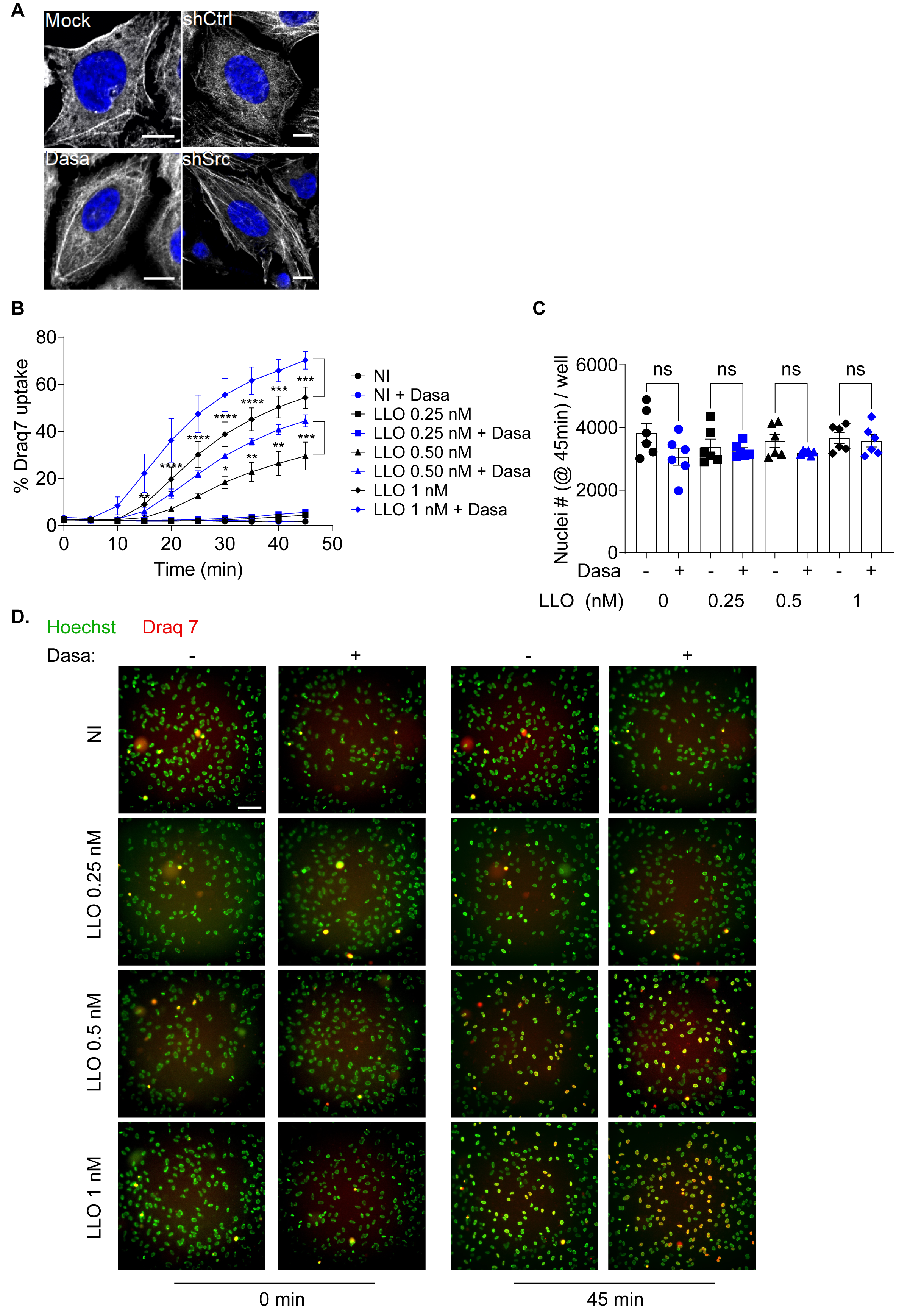

### Fig S4.tif

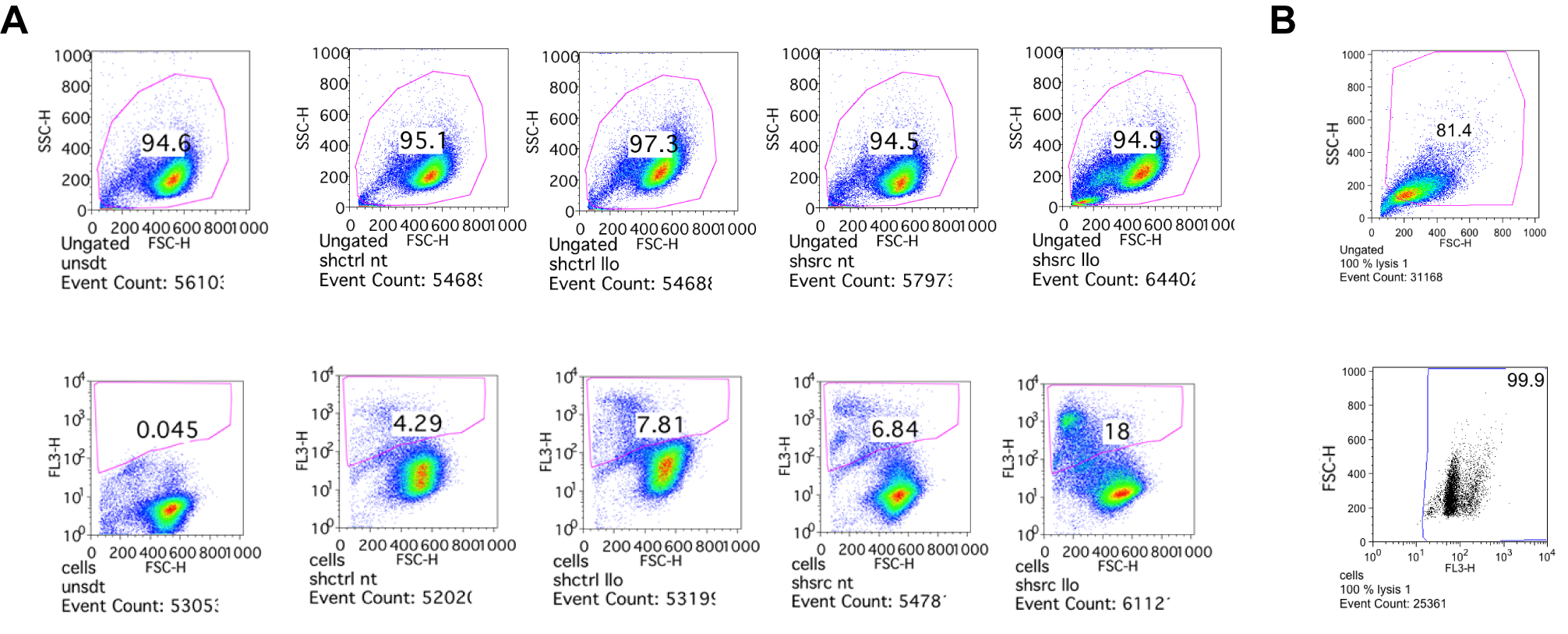

### FigS5 Revised.tif

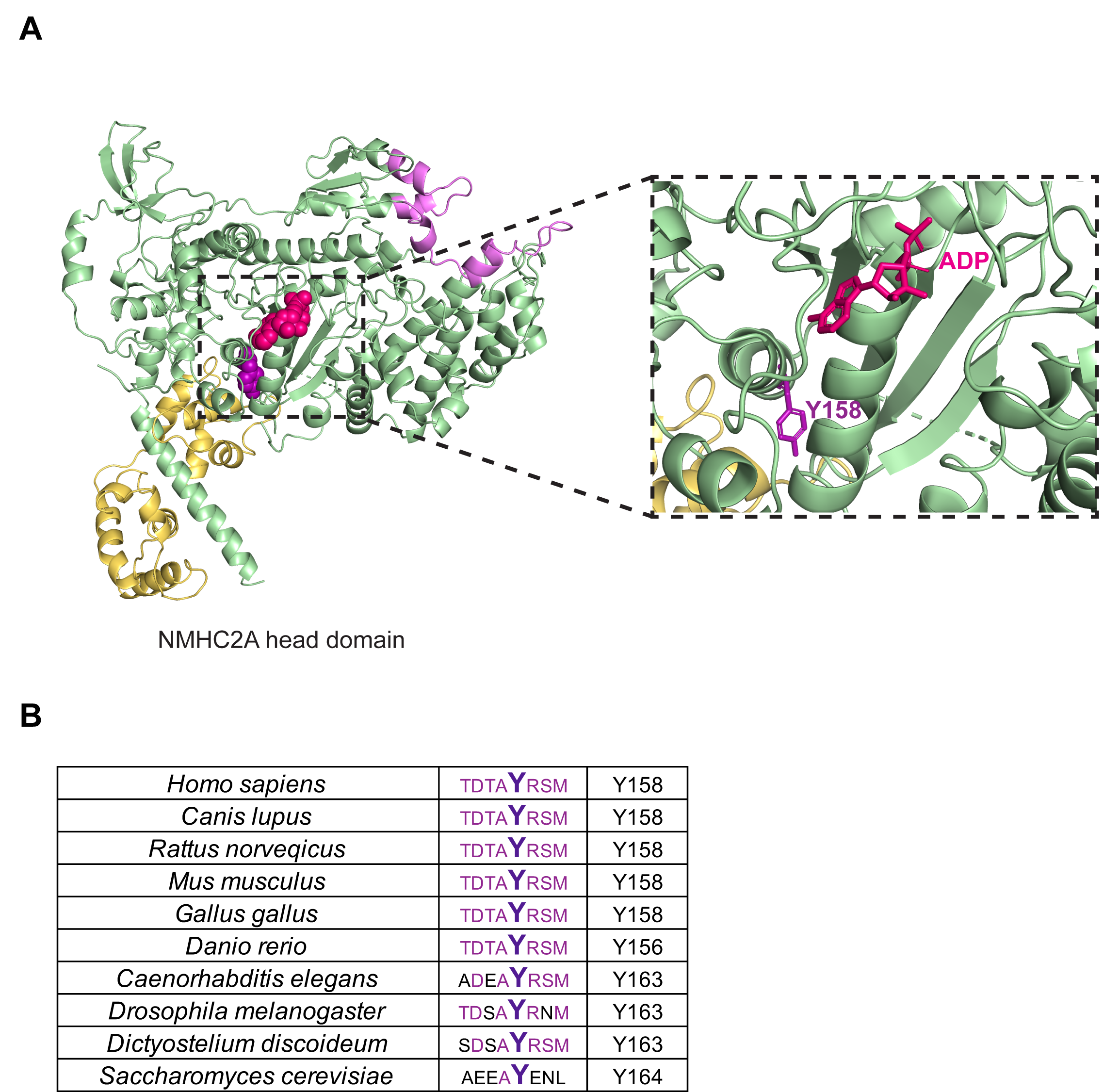

### FigS6 Revised.tif

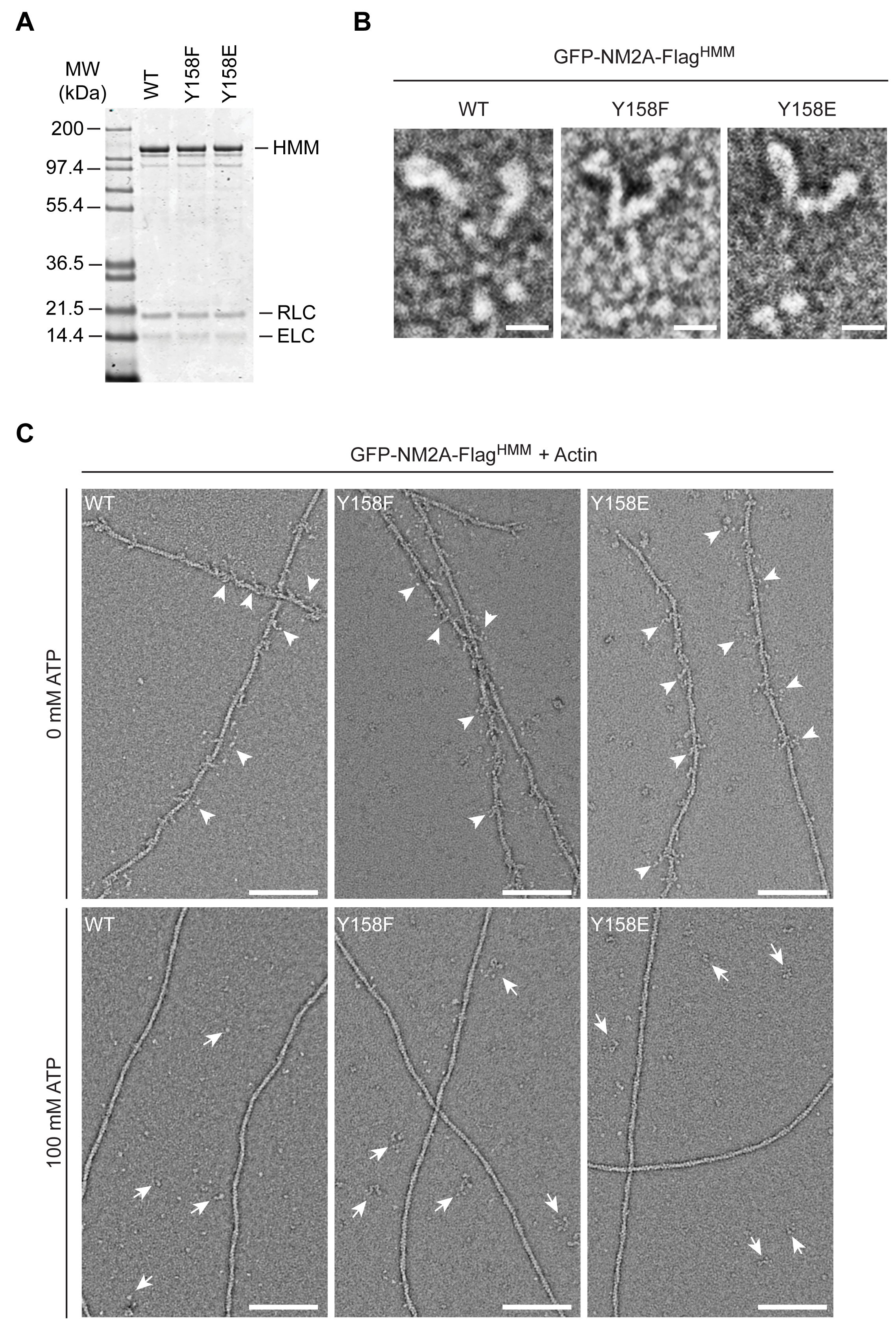

### FigS7 Revised.tif

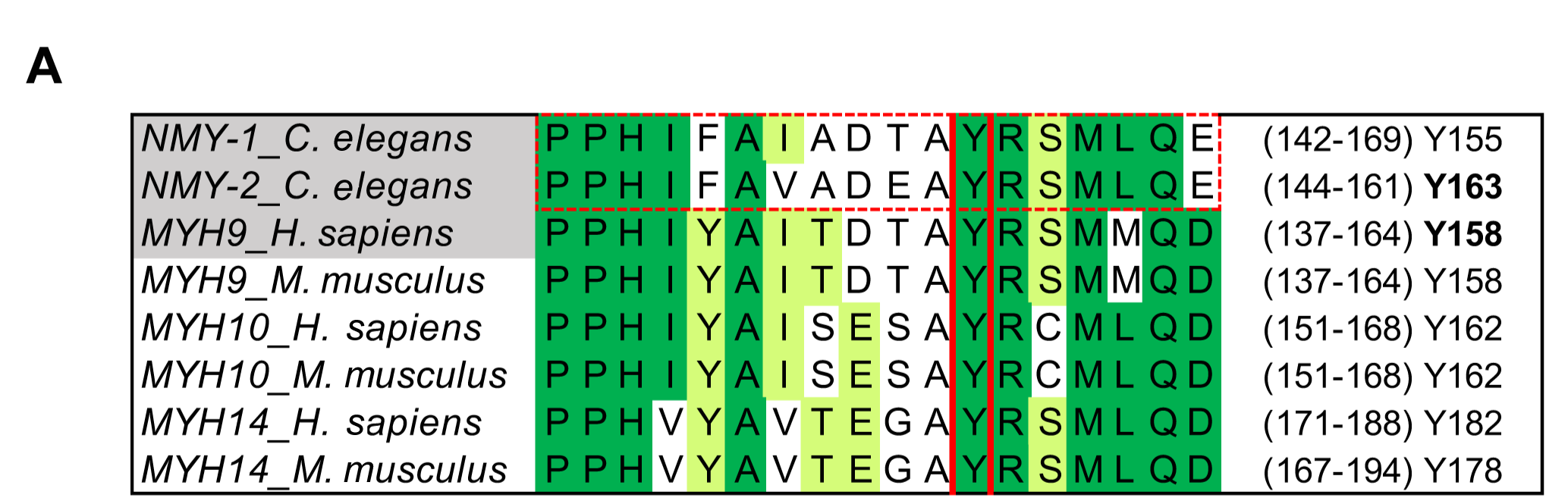
