## Supplementary material for "Src-dependent tyrosine-phosphorylation of NM2A has a protective role against bacterial pore-forming toxins": Table 1

**Table 1.** *C. elegans* strains used in this study.

| Strain | Genotype | Source |
| --- | --- | --- |
| N2 | Ancestral strain | *Caenorhabditis* Genetics Center |
| GCP693 | nmy-2 [prt143(Y163F)]I | This study |
