## Supplementary material for "Src-dependent tyrosine-phosphorylation of NM2A has a protective role against bacterial pore-forming toxins": Table 2

**Table 2.** shRNA sequences for Src knockdown in HeLa cells and single-guide RNAs, single-stranded repair templates and RNAi sequences used in *C. elegans*. Highlighted in grey are the restriction sites used to screen the mutant animals. HindIII for Y163F and Eco47III for Y163E.

| **shRNA Sequences (5’-3’) - HeLa cells** | | **Source** |
| --- | --- | --- |
| *shControl‐SHC016* | CCGGGCGCGATAGCGCTAATAATTTCTCGAGAAATTATTAGCGCTATCGCGCTTTTT | Sigma |
| *shSrc*  *TRCN0000023597* | CCGGGTGGCTTACTACTCCAAACATCTCGAGATGTTTGGAGTAGTAAGCCACTTTTT |  |
| **sgRNAs sequences (5’-3’) - CRISPR in *C. elegans*** | |  |
| *NMY-2_Y163_Guide#1* | TCTTGAAGCATGCTGCGGT*AGG* | Sigma |
| *NMY-2_Y163_Guide#2* | TCGTTCTTGAAGCATGCTG*CGG* |  |
| *NMY-2_Y163_Guide#3* | ATGCTGCGGTAGGCTTCAT*CGG* |  |
| **Single stranded repair templates (5’-3’) - CRISPR in *C. elegans*** | |  |
| *ssODN NMY2_Y163F* | AGA AAG GAG ATG CCA CCA CAC ATA TTT GCT GTT GCT GAT GAA GCT TTC CGC TCA ATG CTC CAG GAA CGA GAT GAT CAG TCA ATT CTC TGC ACG TTA GT | IDT ultramer |
| *ssODN NMY2_Y163E* | AGA AAG GAG ATG CCA CCA CAC ATA TTT GCT GTT GCT GAT GAA GCT GAG CGC TCA ATG CTC CAG GAA CGA GAT GAT CAG TCA ATT CTC TGC ACG TTA GT |  |
| **Diagnosis primer sequences (5’-3’)- CRISPR in *C. elegans*** | |  |
| oAC24 | ACT CTG GCT TGT TCT GCG TT | Sigma |
| oAC564 | GCA CGC GAG ATT TCT CCA G |  |
